# Integration of shoot-derived polypeptide signals by root TGA transcription factors is essential for survival under fluctuating nitrogen environments

**DOI:** 10.1101/2023.10.23.563506

**Authors:** Ryutaro Kobayashi, Yuri Ohkubo, Mai Izumi, Ryosuke Ota, Keiko Yamada, Yoko Hayashi, Yasuko Yamashita, Saki Noda, Mari Ogawa-Ohnishi, Yoshikatsu Matsubayashi

## Abstract

Unlike plants in the field, which experience significant temporal fluctuations in environmental conditions, plants in the laboratory are typically grown in controlled, stable environments. Therefore, signaling pathways evolved for survival in continuously fluctuating environments often remain functionally latent in laboratory settings. Here, we show that TGA1 and TGA4 act as hub transcription factors through which the expression of genes involved in high-affinity nitrate uptake are regulated in response to shoot-derived phloem mobile polypeptides, CEP DOWNSTREAM 1 (CEPD1), CEPD2 and CEPD-like 2 (CEPDL2) as nitrogen (N) deficiency signals, and Glutaredoxin S1 (GrxS1) to GrxS8 as N sufficiency signals. CEPD1/2/CEPDL2 and GrxS1-S8 competitively bind to TGA1/4 in roots, with the former acting as transcription coactivators that enhance the uptake of nitrate, while the latter function as corepressor complexes together with TOPLESS to limit nitrate uptake. *Arabidopsis* plants deficient in TGA1/4 maintain basal nitrate uptake and exhibit growth similar to wild-type plants in a stable N environment, but were impaired in regulation of nitrate acquisition in response to shoot N demand, leading to defective growth under continuously fluctuating N environments where rhizosphere nitrate ions switch periodically between deficient and sufficient states. TGA1/4 are crucial transcription factors that enable plants to survive under fluctuating and challenging N environmental conditions.

## Introduction

Nitrogen (N), which is mostly present as nitrate in soils, is an essential macronutrient that plays a crucial role in plant development. However, due to the high mobility of nitrate ions through soil with water, marked spatiotemporal fluctuations can occur with respect to soil nitrate availability. To cope with such fluctuations in the external N environment, plants have evolved regulatory mechanisms that enable them to modulate the efficiency of root N acquisition in response to their internal N demand and rhizosphere N availability^1, 2^. This systemic adaptive response is mediated by shoot-root communication via phloem-mobile descending CEP DOWNSTREAM 1 (CEPD1), CEPD2 and CEPD-like 2 (CEPDL2) polypeptides^3^. Among these, CEPD1 is induced in shoots in response to the C-TERMINALLY ENCODED PEPTIDE (CEP), which is a root-derived N deficiency signal^4, 5^, while CEPDL2 is induced in response to the shoot’s own N deficiency^6^. CEPD2 is induced in response to both root-derived CEP and shoot N deficiency.

CEPD1/2/CEPDL2 promote nitrate uptake in roots by both transcriptional upregulation of high-affinity nitrate transporter genes, such as *NRT2.1*, and dephosphorylation-mediated post-translational activation of NRT2.1 by the protein phosphatase, CEPD-INDUCED PHOSPHATASE (CEPH), which is also induced by CEPD1/2/CEPDL2^7^. Accordingly, the *cepd1,2 cepdl2* triple mutant showed severely reduced shoot growth, which is characterized by a significant reduction in high-affinity nitrate influx in roots compared with WT.

CEPD1/2/CEPDL2 are non-secreted polypeptides that are composed of 99 to 102 amino acids and belong to a large protein family comprising 21 members in *Arabidopsis*. Currently, these polypeptides are assigned to the plant-specific class III glutaredoxin (Grx) family^8^, although from a structural point of view, there is controversy as to whether individual members are enzymatically active in redox regulation^9^. Of these, ROXY1 and ROXY2 have been shown to regulate petal morphogenesis and microspore formation^10^. Conversely, ROXY18 has been shown to play roles in biotic and abiotic stresses^11, 12^. The Grx family also includes members that are upregulated in shoots under nitrate-sufficient conditions^13–15^, which is opposite to the response associated with CEPD1/2/CEPDL2. Ectopic overexpression of one such gene, *GrxS8*, caused downregulation of nitrate transporter genes accompanied with repression of lateral root development^16^.

There is increasing, but fragmentary, evidence suggesting a possible link among CEPDs, Grxs, TGA family transcription factors, TOPLESS family transcriptional co-repressors^17^ and nitrate signaling. (i) Combining yeast two-hybrid screening data from multiple independent experiments, showed that all 21 members of the class III Grx family are likely to interact with all 10 TGA transcription factors *in vitro*, albeit with varying affinities^11, 18–23^. (ii) Most of the Grxs, except for 4 members in the CEPD1/2/CEPDL2 clade, have been shown to interact with the TOPLESS family of transcriptional co-repressors in the yeast two-hybrid assay^24^. (iii) Co-expression network analysis or transcriptome under varying nitrogen conditions identified TGA1 as a key regulatory factor of the nitrate response^25, 26^. The double mutant of TGA1 and its closest homolog TGA4 exhibited a decrease in lateral root initiation and a reduction in root hair development in response to nitrate treatment^25, 27^. The *tga1 tga4* double mutant, however, showed no observable changes in nitrate uptake activity in roots under normal growth conditions^25^. It has also not yet been definitively established whether TGA1 directly binds to promoters of genes involved in high-affinity nitrate uptake, such as NRT2.1 and CEPH^25, 26^.

To piece together these fragmentary observations, here we analyzed *in planta* signaling network by co-immunoprecipitation-mass spectrometry (CoIP-MS) and chromatin immunoprecipitation followed by sequencing (ChIP-Seq). CEPDs and subset of Grxs competitively bind to TGA1/4, with the former acting as transcription coactivators that enhance the uptake of nitrate, while the latter function as corepressor complexes together with TOPLESS to limit nitrate uptake. *Arabidopsis* plants deficient in TGA1/4 were impaired in demand-dependent regulation of nitrate acquisition, resulting in defective growth under a temporally fluctuating N environment.

## Results

### CEPDL2 acts through interaction with TGA1 and TGA4 transcription factors

We first searched for signaling targets that physically interact with CEPD family polypeptides in roots using an unbiased proteomic approach. Here, GFP-tagged CEPDL2, which has the same signaling properties as untagged CEPDL2^6^, was used as bait for CoIP-MS. Root extracts were prepared from 14-day-old seedlings expressing *GFP-CEPDL2* under the control of its own promoter, and GFP-CEPDL2 was immunoprecipitated from the extracts using immobilized anti-GFP antibodies. Nano liquid chromatography-tandem mass spectrometry (nano-LC-MS/MS) was performed on three replicate immunoprecipitations from *GFP-CEPDL2* seedling roots as well as three control immunoprecipitations from *NRT2.1pro:GFP* seedling roots.

Volcano plots of enrichment and significance showed that TGA1 and TGA4 were the most enriched transcription factors (Fig. 1a). We confirmed direct interaction between CEPDL2 and TGA4 by the yeast two-hybrid assay (Supplementary Fig. 1a). Overexpression of *CEPDL2* in a *tga1,4* double mutant did not upregulate neither of CEPDL2 target genes such as *NRT2.1* and *CEPH* (Fig. 1b, Supplementary Fig. 1b), indicating that *TGA1* and *TGA4* are epistatic to, and act downstream of, *CEPDL2*.

**Fig. 1.**
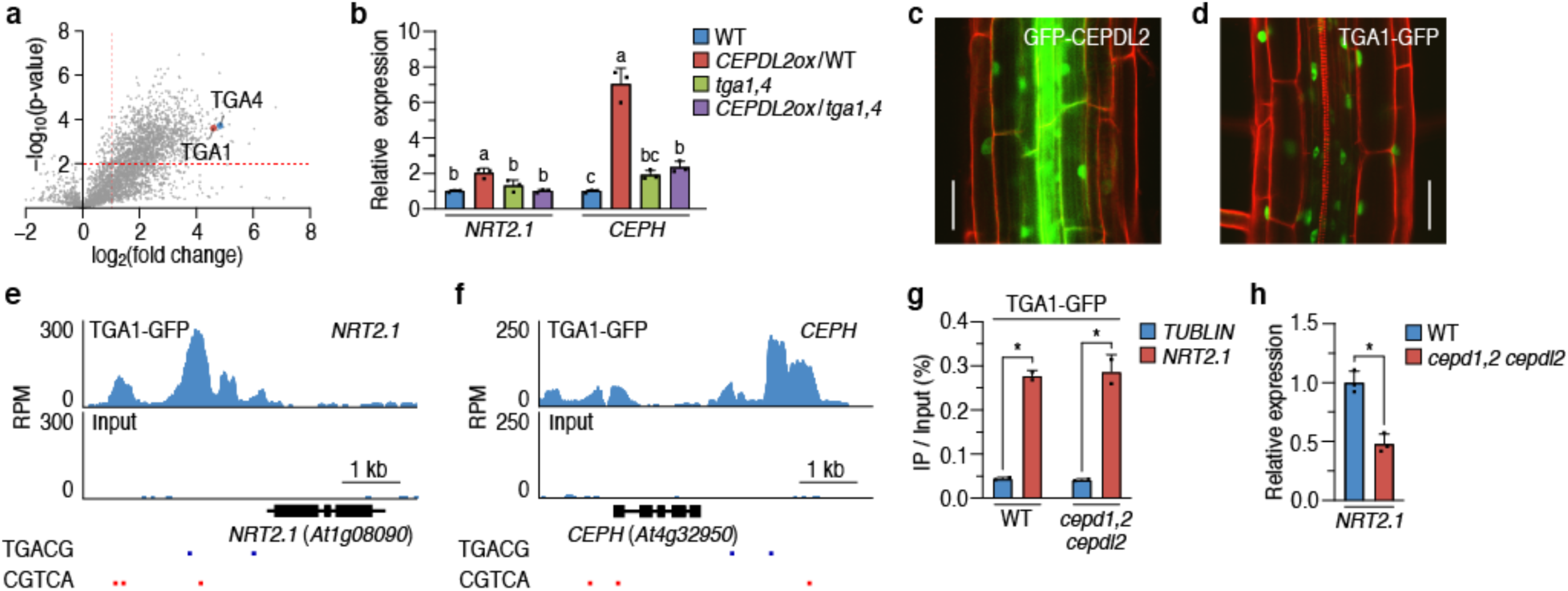
CEPDL2 acts through interactions with TGA1 and TGA4 transcription factors. (**a**) Volcano plot showing the enrichment of proteins co-purified with GFP-CEPDL2 as compared with the GFP control. The Y- and X-axes show p-values and fold changes, respectively. The red dashed line indicates the threshold above which proteins are significantly enriched (p < 0.01, fold change ≥ 2). (**b**) Expression levels of CEPDL2 target genes, *NRT2.1* and *CEPH*, in WT, *CEPDL2ox*/WT, *tga1,4*, and *CEPDL2ox*/*tga1,4* plants grown on 10 mM NO_3_^−^ medium (mean ± SD, P < 0.05 by one-way analysis of variance (ANOVA) with post-hoc Tukey’s test, *n* = 3). (**c-d**) Subcellular localization of GFP-CEPDL2 (c) and TGA1-GFP (d) in roots. Scale bar = 100 µm. (**d-f**) Genome browser views of TGA1 ChIP signals at the *NRT2.1* (e) and *CEPH* (f) promoters, with normalized read counts per million along the Y-axis. Blue and red lines indicate TGACG motifs. (**g**) ChIP-qPCR analyses of TGA1-GFP binding to the *NRT2.1* promoter in roots of WT and *cepd1,2 cepdl2* plants (mean ± SD, *P < 0.05 by two-tailed non-paired Student’s t test, *n* = 2). (**h**) Expression levels of *NRT2.1* in the same plants, as in (**g**) (mean ± SD, *P < 0.05, *n* = 3).

### CEPDs are required for TGA-mediated transcriptional activation, but not for promoter binding of TGA

To determine the binding sites of TGA1 across the genome *in vivo*, we used chromatin immunoprecipitation followed by sequencing (ChIP-Seq). For the ChIP-Seq analysis, we used transgenic *Arabidopsis* plants expressing TGA1-GFP using their own promoter, with plants expressing GFP (*NRT2.1pro:GFP*) as a negative control. TGA1-GFP localized in the nucleus of endodermal, cortical, and epidermal cell layers in roots, which overlaps with the localization of CEPDL2 (Fig. 1c, d). We performed ChIP-Seq on 16-day-old roots and identified 1,105 TGA1 *in vivo* target loci (q-value < 10^-40^) from the TGA1-GFP-expressing plants that were not identified in a GFP control line (Supplementary Dataset 1). The TGA1 targets included genes related to high-affinity nitrate uptake, such as *NRT2.1* and *CEPH* (Fig. 1e, f), while *NRT1.1*, which is involved in low-affinity nitrate uptake, was not detected. We confirmed binding motifs for TGA1 enriched in the ChIP-Seq reads as “TGACG” (e-value = 4.0 × 10^-7^) and the most enriched GO term as “response to nitrate” (Supplementary Fig. 1c, d). We further examined the overlap between the 296 differentially expressed genes (DEGs) (fold-change ≥ 2) induced in *35S:CEPDL2* plant roots^6^ and the TGA1 ChIP-Seq target genes. We identified 48 genes that overlapped (Supplementary Dataset 2), including *NRT2.1*, *NRT2.2*, *NRT3.1*, *NRT1.5*, and *CEPH*, all of which have been reported to play key roles in nitrate acquisition.

We also tested whether CEPD1/2/CEPDL2 are required for the binding of TGA1 to target gene promoters. We produced transgenic *Arabidopsis* plants expressing TGA1-GFP using the own promoter in WT plants and the *cepd1,2 cepdl2* triple mutant. ChIP-qPCR analysis showed that TGA1 binds to the promoter of *NRT2.1* in the *cepd1,2 cepdl2* triple mutant with a comparable affinity in WT, even though the expression level of *NRT2.1* declined to 48.3% of that in WT (Fig. 1g, h). These results indicate that CEPD1/2/CEPDL2 are required as coactivators for TGA-mediated transcriptional activation of the target genes, but that they are not required for the binding of TGA to the target-binding regions.

### Loss of *TGA1/4* counteracted the growth defects observed in the *cepd1,2 cepdl2* triple mutant

We previously reported that the *cepd1,2 cepdl2* triple mutant shows severe growth defects, such as a decrease in shoot fresh weight and reduced expression of the *NRT2.1* and *CEPH* genes^6^. However, it has been reported that the *tga1,4* double mutant shows no such growth defects, which we also confirmed in this study (Fig. 2a-c). To test the genetic interactions between *CEPD*s and *TGA*s, we crossed the *cepd1,2 cepdl2* triple mutant with the *tga1,4* double mutant. Unexpectedly, the introduction of the *tga1,4* mutation counteracted the growth defects in the phenotypes of the *cepd1,2 cepdl2* triple mutant, restoring them to WT levels (Fig. 2a-c). These results indicate that TGA1/4 have a positive effect on regulating the transcription of nitrate uptake genes after binding to coactivator CEPD1/2/CEPDL2 but exhibits a negative effect on the transcription of nitrate uptake genes in the absence of CEPDs. In the *tga1,4* mutant, both positive and negative effects are lost.

**Fig. 2.**
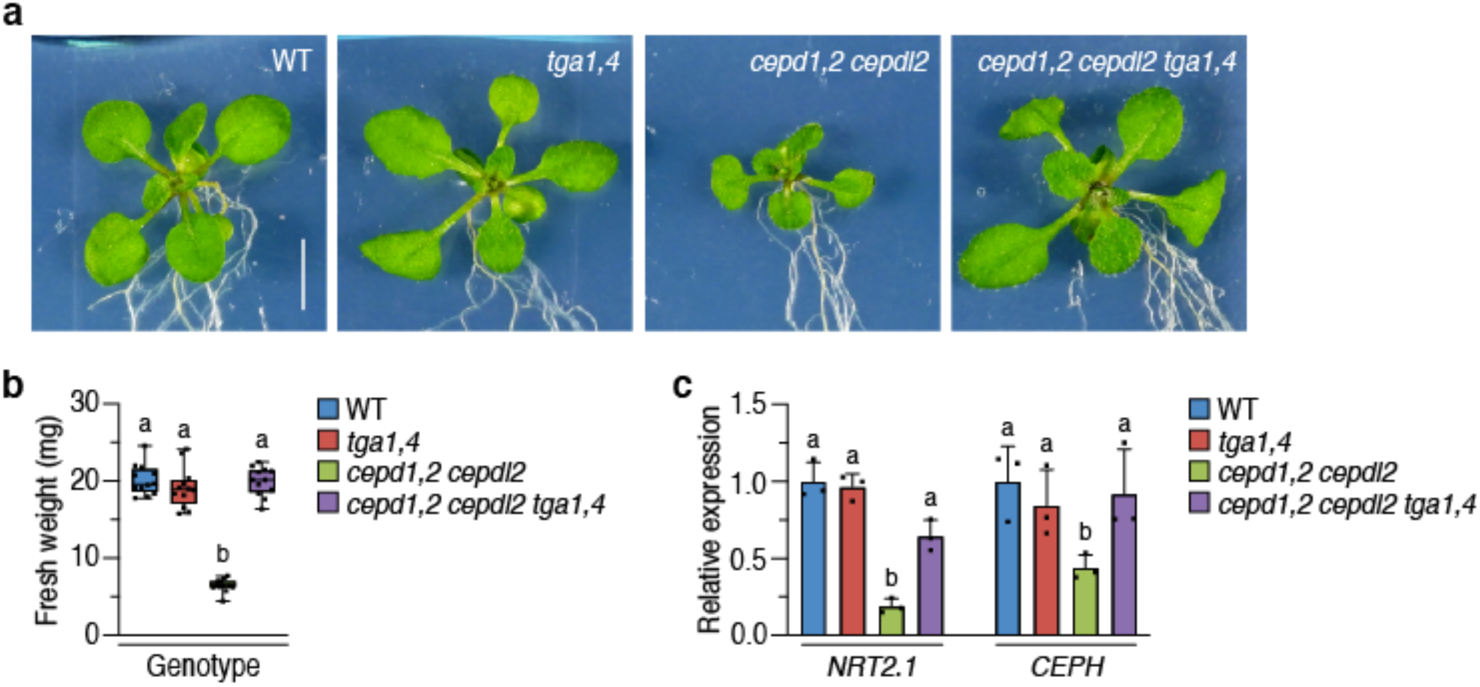
Loss of *TGA1/4* counteracted the growth defects of the *cepd1,2 cepdl2* triple mutant. (**a**) Phenotypes of 14-day-old WT, *tga1,4*, *cepd1,2 cepdl2*, and *cepd1,2 cepdl2 tga1,4* plants grown on 3 mM NO_3_^−^ medium. Scale bar = 5 mm. (**b**) Fresh weight of 14-day-old shoots of WT, *tga1,4*, *cepd1,2 cepdl2*, and *cepd1,2 cepdl2 tga1,4* plants grown on 3 mM NO_3_^−^ medium (mean ± SD, P < 0.05, *n* = 12). (**c**) Expression levels of *NRT2.1* and *CEPH* genes in WT, *tga1,4*, *cepd1,2 cepdl2*, and *cepd1,2 cepdl2 tga1,4* plants grown on 3 mM NO_3_^−^ medium (mean ± SD, P < 0.05, *n* = 3).

### GrxS1-S8 family polypeptides have roles opposite to those of CEPDs

When considering the suppressive effect of TGA1/4 on the transcription of nitrate uptake genes in the absence of the coactivator CEPDs, we considered that a corepressor that interacts with TGA1/4 could provide a plausible explanation for this suppression of transcription. We focused on the GrxS1/S2/S3/S4/S5/S6/S7/S8 family polypeptides as potential candidates because, despite having a sequence that is highly similar to CEPDs, *GrxS8* has been reported to suppress the transcription of nitrate transporter genes when overexpressed^16^.

*GrxS1* to *GrxS8* genes are expressed almost exclusively in the phloem tissues of leaves (Fig. 3a, Supplementary Fig. 2a, b) and upregulated in leaves under nitrate-sufficient conditions, a response that is opposite to that observed with CEPDs (Supplementary Fig. 2c). When a *GFP-GrxS5* construct driven by the native *GrxS5* promoter was introduced into WT plants, we observed GFP-GrxS5 signals in the nucleus of endodermal, cortical, and epidermal cell layers in roots (Fig. 3b), indicating that the GrxS5 functions as a phloem-mobile shoot-to-root signal. This GFP-GrxS5 transgene, which caused over-accumulation of *GrxS5* transcripts, suppressed the expression of 415 genes, including both high- and low-affinity nitrate uptake genes (Fig. 3c, Supplementary Fig. 2d, e, Supplementary Dataset 3). Notably, among the genes that were downregulated by GrxS5, 11 genes, including *NRT2.1*, *NRT2.2*, *NRT3.1*, *NRT1.5*, and *CEPH*, were direct targets regulated by the CEPD/TGA pathway (Supplementary Table 1).

**Fig. 3.**
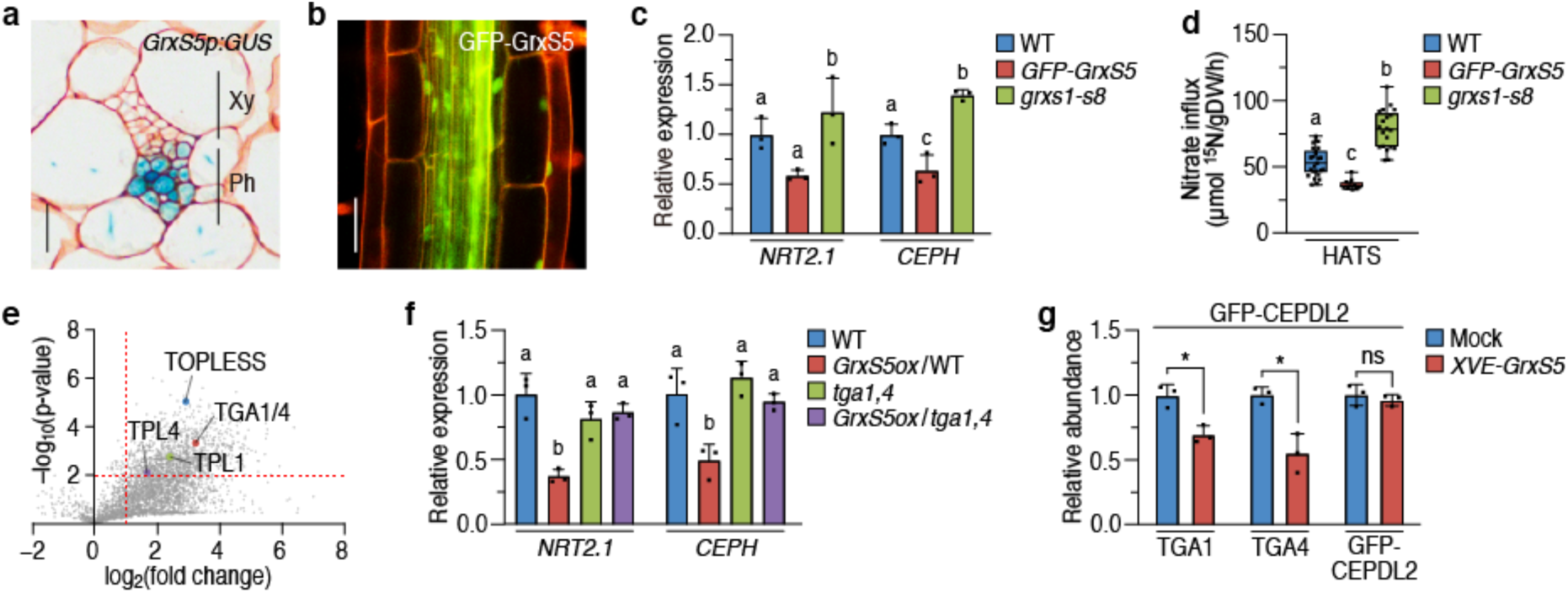
GrxS1-S8 family polypeptides play roles opposite to those of CEPDs and compete with CEPDs to interact with TGA1/4. (**a**) Histochemical localization of GUS activity in cross section of the leaf vascular tissues expressing *GrxS5pro:GUS* gene. Abbreviations: Xy, xylem; Ph, phloem. Scale bar = 20 µm. (**b**) Detection of GFP-GrxS5 signals in the root of 9-day-old plants expressing *GrxS5pro:GFP-GrxS5*. Scale bar = 100 µm. (**c**) Expression levels of nitrate uptake genes in WT, *GFP-GrxS5*, and *grxs1-s8* plants grown on 10 mM NO_3_^−^ medium (mean ± SD, P < 0.05, *n* = 3). (**d**) Activity of high-affinity nitrate uptake system (HATS) of 14-day-old WT, *GFP-GrxS5* and *grxs1-s8* plants grown on 10 mM NO_3_^−^ medium (mean ± SD, P < 0.05, *n* = 10-25). (**e**) Volcano plot showing the enrichment of proteins co-purified with GFP-GrxS5 as compared with GFP control. (**f**) Expression levels of GrxS5-target genes, *NRT2.1* and *CEPH*, in WT, *GrxS5ox*/WT, *tga1,4* and *GrxS5ox*/*tga1,4* plants grown on 10 mM NO_3_^−^ medium (mean ± SD, P < 0.05, *n* = 3). (**g**) Relative abundance of TGA1 and TGA4 co-immunoprecipitated with GFP-CEPDL2 in GrxS5 overexpressing plants (mean ± SD, *P < 0.05, *n* = 3).

When ectopically overexpressed, all genes from *GrxS1* to *GrxS8* suppressed the expression of *NRT2.1* (Supplementary Fig. 2f). Conversely, in the octuple mutant *grxs1-s8* (i.e., *grxs1-1*, *grxs2-1*, *grxs3-1*, *grxs4-1*, *grxs5-1*, *grxs6-1*, *grxs7-1*, *grxs8-1*) generated by T-DNA and CRISPR/Cas9-mediated mutation (Supplementary Fig. 2g), the mutations had positive effects on the expression levels of nitrate uptake genes (Fig. 3c, Supplementary Fig. 2e), increasing both high- and low-affinity nitrate uptake activities (Fig. 3d, Supplementary Fig. 2h). This *grxs1-s8* octuple mutant showed a decrease in survival rate under excess nitrate (120 mM) conditions (Supplementary Fig. 2i), likely due to the lack of negative feedback on excess nitrate uptake. These findings indicate that while CEPD1/2/CEPDL2 and GrxS1-S8 function as phloem-mobile shoot-to-root signals, they play opposite roles in regulating nitrate acquisition.

### CEPDL2 and GrxS5 competitively interact with TGA1/4

We searched for signaling targets that physically interact with the GrxS1-S8 family of polypeptides in roots by CoIP-MS using GFP-GrxS5 as bait. Co-IP data showed the interactions between GFP-GrxS5 and TGA1/4 (conserved sequences between TGA1 and TGA4 were detected), in addition to TOPLESS, TOPLESS-related 1 (TPL1), and TPL4 (Fig. 3e), which suggests that GrxS5 interacts with TOPLESS family corepressors to inhibit TGA-mediated transcription. We confirmed direct interaction between GrxS5 and TGA4 by the yeast two-hybrid assay (Supplementary Fig. 3a). However, based on their fold-change values, the binding affinity of GrxS5 to TGA1/4 appears to be weaker than that of CEPDL2 to TGA1/4.

Overexpression of *GrxS5* in a *tga1,4* double mutant background failed to downregulate *NRT2.1* and *CEPH*, indicating that *TGA1* and *TGA4* also acts downstream of *GrxS5* (Fig. 3f, Supplementary Fig. 3b). In contrast, loss of *TGA1/4* had no effect on GrxS5-dependent downregulation of *NRT1.1*, suggesting that GrxS5 represses low affinity nitrate uptake in a pathway independent of the TGA1/4 (Supplementary Fig. 3c).

We hypothesized that, due to their striking sequence similarity, might bind to TGA in a competitive manner. To test this possibility, we overexpressed GrxS5 using an estradiol-inducible system in plants expressing GFP-CEPDL2 and immunoprecipitated the GFP-CEPDL2/TGA complex using an anti-GFP antibody. CoIP-MS data showed that, under conditions where GFP-CEPDL2 was immunoprecipitated with comparable efficiency, the amount of co-immunoprecipitated TGA1 and TGA4 decreased in GrxS5-overexpressing plants compared to that in mock plants (Fig. 3g, Supplementary Fig. 3d), indicating that GrxS5 competes with CEPDL2 for binding to both TGA1 and TGA4. Based on these results, we propose a model in which CEPD1/2/CEPDL2 and GrxS1-S8 competitively bind to TGA1/4, with the former acting as transcription coactivators that enhance the uptake of nitrate, while the latter function as corepressor complexes together with TOPLESS to limit nitrate uptake.

### TGA1/4 are critical for plant survival under a temporally fluctuating N environment

Despite the presumed mechanistic importance of TGA1/4 in N acquisition, the *tga1,4* double mutant did not exhibit any visible phenotypes under either N sufficient (3 mM NO_3_^−^) (Fig. 2a) or low N (0.1 mM NO_3_^−^) conditions (Fig. 4a). These results raise the question of how a system that integrates N deficiency signals and N sufficiency signals from aboveground parts through root TGA1/4 transcription factors contributes to plant survival. Since this long-range signaling system should regulate root nitrate uptake in response to aboveground N demand, we hypothesize that TGA1/4 are particularly important in environments where shoot N demand and rhizosphere N availability fluctuate over time.

**Fig. 4.**
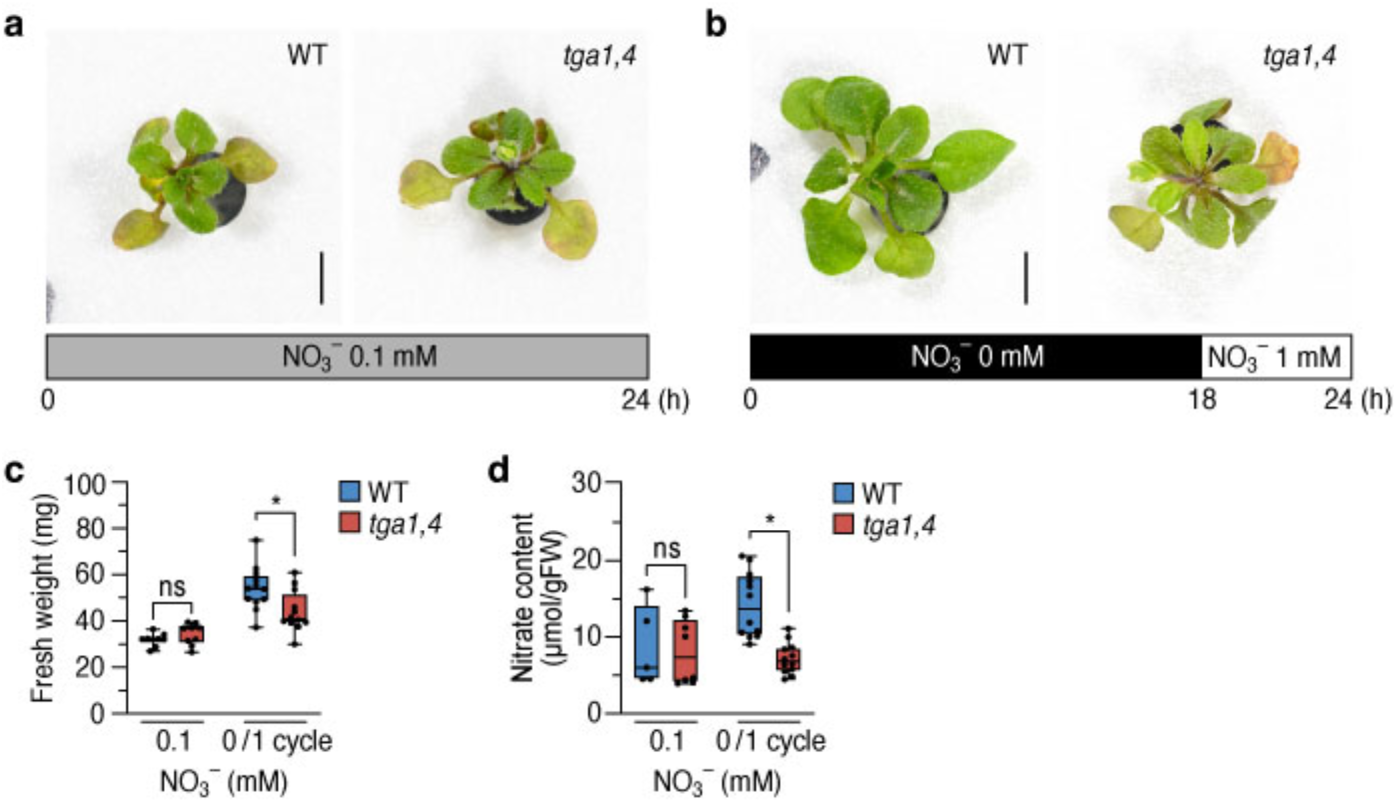
TGA1/4 are critical for plant survival under temporally fluctuating nitrogen environment. (**a**) Phenotypes of WT and *tga1,4* plants cultured for 7 d under continuous low nitrogen condition (0.1 mM NO_3_^−^) Scale bar = 5 mm. (**b**) Phenotypes of WT and *tga1,4* plants cultured for 7 d under fluctuating nitrogen conditions consisting of 18 h of nitrogen-deficient conditions (0 mM NO_3_^−^) followed by 6 h of nitrogen-sufficient conditions (1 mM NO_3_^−^) for the duration of the experiment. Scale bar = 5 mm. (**c**) Fresh weight of WT and *tga1,4* plants grown for 7 d under continuous low nitrogen or fluctuating nitrogen conditions (mean ± SD, *P < 0.05, *n* = 9-12). (**d**) Nitrate content in the shoots of WT and *tga1,4* plants grown for 7 d under continuous low nitrogen or fluctuating nitrogen conditions (mean ± SD, *P < 0.05, *n* = 5-12).

To investigate this possibility, we cultured WT and *tga1,4* mutant plants in a hydroponic system in which N-deficient (0 mM NO_3_^−^) and N-sufficient (1 mM NO_3_^−^) conditions were alternated for 18 h of and 6 h of each day over 7 d, respectively. Under this fluctuating N environment, a clear difference in phenotypes between the WT and *tga1,4* mutant plants was observed, with the *tga1,4* mutant showing stunted growth characterized by smaller rosettes and reduced shoot biomass compared with WT plants (Fig. 4b, c). The *tga1,4* mutant also exhibited a significant reduction in nitrate content in shoots (Fig. 4d), accompanied by early leaf yellowing that is typically induced by N deprivation. These results indicate that TGA1/4 transcription factors are indispensable for plant survival under temporally fluctuating N conditions.

### Demand-dependent regulation of nitrate acquisition was impaired in *tga1,4* mutants

Finally, we explored the reasons why the *tga1,4* mutant failed to adapt to fluctuating N environments by analyzing the time-course expression levels of N uptake-related genes during repetitive cycles of N deficiency and sufficiency. Since the nitrate content in the shoots of the *tga1,4* mutant was already lower than that of the WT at 72 h after the beginning of the repetitive cycles (Supplementary Fig. 3e), we examined the expression of *CEPDs* and *GrxS5* in leaves, as well as nitrate uptake genes in roots from 24 h to 72 h using RT-qPCR.

In shoots of both the *tga1,4* mutant and the WT, the expression of *CEPDL2*, *CEPD2*, and *CEPD1* were induced under N deficient-conditions and repressed under N-sufficient conditions (Fig. 5a). Conversely, in both the *tga1,4* mutant and WT plants, the expression of *GrxS5* was repressed under N-deficient conditions and induced under N-sufficient conditions (Fig. 5a).

**Fig. 5.**
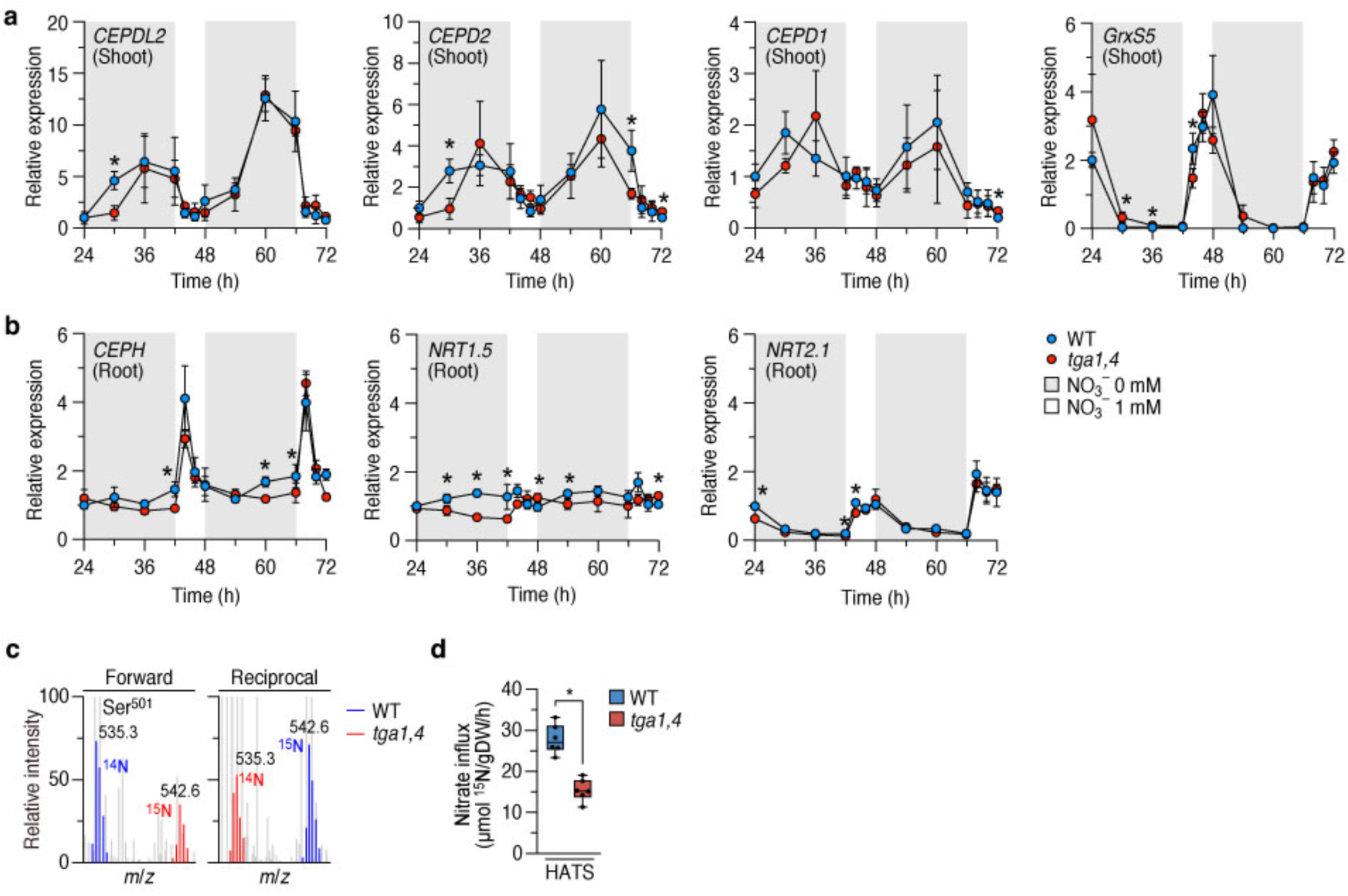
The *tga1,4* mutant was impaired in demand-dependent regulation of nitrate acquisition. (**a**) Expression levels of *CEPDL2*, *CEPD2*, *CEPD1* and *GrxS5* in leaves of WT and *tga1,4* plants fluctuating nitrogen conditions consisting of 18 h of nitrogen-deficient conditions (0 mM NO_3_^−^) followed by 6 h of nitrogen-sufficient conditions (1 mM NO_3_^−^) (mean ± SD, *P < 0.05, *n* = 3). Sampling was performed at 2-6 h intervals from 24 h after the beginning of culture till 72 h. (**b**) Expression levels of *CEPH*, *NRT1.5* and *NRT2.1* in WT and *tga1,4* roots under fluctuating nitrogen condition (mean ± SD, *P < 0.05, *n* = 3). (**c**) MS-based quantification of Ser^501^ dephosphorylated active form of NRT2.1 in WT and *tga1,4* plants at 66 h after the beginning of culture under fluctuating nitrogen condition. Mass spectra of the NRT2.1 Ser^501^ dephosphorylated peptide (position 496-509 in NRT2.1) derived from roots of the plants metabolically labeled by ^14^N WT/^15^N *tga1,4* (forward) and ^15^N WT/^14^N *tga1,4* (reciprocal) combinations were shown. (**d**) Activity of high-affinity nitrate uptake system (HATS) of WT and *tga1,4* plants at 66 h (mean ± SD, P < 0.05, *n* = 6).

In roots, during N-deficient conditions, the induction of CEPDL2 and CEPD1/2 in leaves and their subsequent translocation to the roots led to gradual upregulation of *CEPH* and *NRT1.5* at the late stage of nitrogen deficient periods in WT plants (Fig. 5b). However, such N demand-dependent responses were not observed in the roots of the *tga1,4* mutant. The transcription of *NRT2.1* was strongly repressed during N-deficient conditions and recovered to the basal levels during N-sufficient conditions in both the *tga1,4* mutant and the WT plants (Fig. 5b), which corresponds to the nitrate-inducible nature of *NRT2.1* and inability of CEPDs to induce *NRT2.1* under N-deficient conditions^3^. During N-sufficient conditions, *CEPH* also showed rapid induction in both WT and *tga1,4* plants, possibly reflecting its cell-autonomous nitrate-inducible nature.

Using ^15^N-metabolic labeled quantitative phosphoproteomic profiling, we further analyzed the phosphorylation level of Ser^501^ of NRT2.1, a residue that functions as a negative phospho-switch. The WT and *tga1,4* plants were cultured on medium containing ^14^NO_3_^−^ or ^15^NO_3_^−^ in a reciprocal manner for biological duplicates (^14^N WT/^15^N *tga1,4* [forward experiment] and ^15^N WT/^14^N *tga1,4* [reciprocal experiment]). At 66 h after the beginning of the repetitive cycles, which corresponds to the end of the third N-deficient period, the Ser^501^–dephosphorylated active form of NRT2.1 in the *tga1,4* mutant decreased to 48% and 74% of that in WT plants in the forward and reciprocal experiments, respectively, without any alteration in NRT2.1 protein expression levels (Fig. 5c, Supplementary Fig. 3f). Consequently, *tga1,4* plants exhibited only 55% of the high affinity nitrate uptake activity of WT plants at 66 h (Fig. 5d). These findings suggest that the inability of *tga1,4* mutants to adapt to fluctuating N environments is due to a decrease in both CEPH-mediated activation (dephosphorylation) of NRT2.1 high affinity nitrate uptake and the NRT1.5-mediated translocation of nitrate ions from roots to shoots.

## Discussion

Our findings elucidated the crucial roles played by TGA1/4 transcription factors under fluctuating N conditions. These roles have been overlooked because *tga1,4* mutants do not show distinct phenotypes under either continuous N-deficient or continuous N-sufficient conditions, despite their presumed involvement in N responses. Under continuous N-deficient conditions, N demand signals derived from leaves are strongly upregulated. However, due to the absence of rhizospheric N, long-distance signaling through CEPD1/2/CEPDL2 and TGA1/4 does not exert its intrinsic effect. The environmental conditions under which the CEPD-TGA system is most effective are when N availability is temporally fluctuating, i.e., where shoots remain in a N-deficient state while nitrate becomes available again in the rhizosphere.

TGA1/4 transcription factors not only interact with the shoot-derived N demand signals CEPD1/2/CEPDL2, but also integrate the N sufficiency signals GrxS1 to GrxS8 in order to optimize the expression of nitrate uptake-related genes in roots. The *grxs1-s8* octuple mutant generated in this study showed a decrease in survival rate under excess nitrate conditions.

However, since such high concentrations of nitrate ions are rare in nature, the GrxS1-S8 pathway is likely a feedback mechanism that prevents the CEPD1/2/CEPDL2 pathway from becoming overactive which leads to uptake of more nitrate than needed, rather than an adaptation to extreme environments. In fact, the binding of GrxS5 to TGA1/4 was considerably weaker compared to CEPDL2. Therefore, the function of TGA1/4, which is suppressed by GrxS1-S8 during N sufficiency, would be switched rapidly to the transcriptional activation of high-affinity nitrate uptake genes by competitive binding with CEPD1/2/CEPDL2.

Nitrate in soils is an essential macronutrient that plays a crucial role throughout plant development; however, due to its high mobility through soil with water, soil nitrate availability fluctuates over time. Plants take up dissolved nitrate ions in the soil pore water, which is the water located between the soil particles. Real-time measurement of the nitrate concentration in the soil pore water revealed rapid changes in nitrate concentration following irrigation or precipitation cycles on a timescale of hours to days^28^. The rapid increase in soil water content leads to a temporary decrease in nitrate concentration in the pore water. In contrast, as the soil water content decreases, the nitrate concentration in the pore water increases again due to the re-diffusion of nitrate from soil particles.

It is also known that water-deficient conditions significantly reduce the nitrate content in roots^29^. A marked reduction in soil water content hinders the nitrate uptake due to reduced root surface area in contact with soil pore water. Thus, nitrate uptake is significantly influenced by the constant fluctuations of the water content in the soil. Rain is always falling on the land somewhere, increasing the water content in the soil. Conversely, insolation causes the moisture in the soil to evaporate, decreasing the water content. In such a temporally variable environment, the CEPD-TGA system plays an important role in maintaining adequate nitrate uptake to meet N demand of the aboveground tissues. Plants, just like animals, would likely be unable to survive in the challenging natural environment without binge eating in response to hunger.

## Methods

### Growth conditions

*Arabidopsis* ecotype Col or Nössen plants were used as the WT control depending on the mutant ecotype. Surface-sterilized seeds were sown on B5 medium (1% sucrose) solidified using 1.5% agar in 13 × 10 cm plastic plates (18 seeds × 3 rows/plate) and grown vertically at 22°C with continuous light at an intensity of 80 µmol·m^−2^·s^−1^. After 7 d, seedlings were transferred to modified Murashige-Skoog medium solidified using 1.5% agar in 13 × 10 cm plastic plates (12 seedlings/plate) and further grown vertically at 22°C with continuous light. Modified Murashige-Skoog medium for 10 mM NO_3_^−^ medium, 10 mM KNO_3_ was added as the sole source of N and K, and half-strength concentrations of the other elements and 0.5% sucrose, adjusted to pH 5.7 with KOH. For 3 mM NO_3_^−^ medium, 3 mM KNO_3_ and 7 mM KCl were added. For 1 mM NO_3_^−^ medium, 1 mM KNO_3_ and 9 mM KCl were added. For 0.1 mM NO_3_^−^ medium, 0.1 mM KNO_3_ and 9.9 mM KCl were added. For N-depleted medium, 10mM KCl was added solely as the K source. For 50 mM or 120 mM NO_3_^−^ medium, 50mM or 120 mM KNO_3_ was added solely as the N source, respectively.

### Mutant plants

The *tga1,4* double mutants were obtained from the SALK T-DNA mutant collection (*tga1-1*, SALK_028212; *tga4-1*, SALK_127923). This *tga1,4* (Col) mutant was backcrossed three times to wild-type Nössen to generate *tga1,4* (Nos) mutant. The *cepd1-1 cepd2-1 cepdl2-1* triple mutants (Nössen background) were described previously. The *grxs1* mutant was obtained from the SALK T-DNA mutant collection (SALK_032946). The *grxs2* mutant, the *grxs6* mutant and the *grxs3/4/5/7/8* quintuple mutants were generated using a CRISPR/Cas9 system. The U6.26 promoter and guide RNA sequences were cloned into pKIR1.0^30^ and transformed into Col-0. We screened transformants by PCR and sequencing and found a 11-bp deletion accompanied with a 1-bp insertion in *GrxS2*, and a 13-bp deletion in *GrxS6*. We also found that the *grxs3/4/5/7/8* quintuple mutants have a 1-bp insertion in *GrxS3,* a 6650-bp deletion in *GrxS4/5/7* (At1g15670-At1g15690), and a 6-bp deletion accompanied with a 2-bp insertion in *GrxS8*. The CRISPR/Cas9 construct was then removed to ensure genetic stability.

### Transgenic plants

For co-immunoprecipitation using GFP-GrxS5, we amplified the 2-kb 5’-upstream region of the *GrxS5* gene, the cDNA fragment of *GrxS5* coding region and the cDNA fragment of GFP using PCR. These three fragments were cloned by translational fusion using four-component ligation into the binary vector pCAMBIA1300-BASTA using NEBuilder HiFi DNA Assembly Kit. The resulting *GFP-GrxS5* construct was introduced into wild-type Col. For expression of GFP as a positive control of co-immunoprecipitation, we amplified the 2-kb 5’-upstream region of the *NRT2.1* gene, the cDNA fragment of GFP using PCR. These two fragments were cloned by translational fusion using three-component ligation into the binary vector pCAMBIA1300-BASTA. The resulting *proNRT2.1:GFP* construct was introduced into wild-type Nössen. The *GFP-CEPDL2*/*cepd1,2 cepdl2* plants were described previously. For overexpression analysis of *CEPDL2* and *GrxS5* in the *tga1,4* mutants, we obtained the open reading frames of each gene by RT–PCR with complementary DNA from the cotyledons, and cloned them downstream of the CaMV 35S promoter in pCAMBIA1300-BASTA. The constructs were introduced into wild-type Col or *tga1,4* (Col) mutant. For analysis of TGA1-GFP expression, the full-length *TGA1* genomic fragment containing the 2-kb promoter region and the GFP-coding region were ligated in-frame in this order into the binary vector pCAMBIA1300-BASTA. The resulting *TGA1-GFP* construct was introduced into wild-type Nössen and *cepd1,2 cepdl2* triple mutant. For overexpression analysis of *GrxS1-S8* in wild-type Col, we obtained the open reading frames of each gene by RT–PCR with complementary DNA from the cotyledons, and cloned them downstream of the CaMV 35S promoter in pBI121. For estradiol-inducible expression of *GrxS5* in *GFP-CEPDL2* plants, a cDNA of *GrxS5* was obtained by RT-PCR using total RNA isolated from *Arabidopsis* leaves. The resulting fragment was cloned into the estradiol-inducible binary vector pER8^31^. The construct was introduced into wild-type Col. For estradiol-mediated *GrxS5* expression, roots of 14-day-old transgenic plants expressing *GrxS5* under control of the estradiol-inducible promoter were treated with β-estradiol for 18 h by direct addition of 2 ml of 50 μM β-estradiol solution onto the transgenic plants on the plates.

### Co-immunoprecipitation

Surface-sterilized seeds of *GFP-CEPDL2*, *GFP-GrxS5*, *proNRT2.1:GFP* plants were sown on B5 medium (1% sucrose) solidified using 1.2% agar in 13 × 10 cm plastic plates (18 seeds × 3 rows/plate) and grown vertically at 22 °C with continuous light at an intensity of 80 µmol·m^−2^·s^−1^. After 7 d, the seedlings were transferred to 10 mM NO_3_^−^ medium and further grown vertically at 22 °C with continuous light. Roots of 14-day-old plants (≈500 mg) were frozen in liquid nitrogen and ground to a fine powder using a Multi-beads shocker (Yasui Kikai Corp., Japan). Total proteins were extracted by mixing 500 mg of ground tissue with 2.0 ml of extraction buffer (50 mM Tris-HCl, 150 mM NaCl, 1% Triton X-100, cOmplete Mini EDTA-free Protease Inhibitor Cocktail [Roche, 1 tablet per 10 ml]), followed by brief sonication in an ultrasonic bath for 5 s. After incubation on ice for 30 min, the homogenate was centrifuged for 15 min at 10,000 × *g* and 4°C, and then the supernatant was collected in a new tube. Anti-GFP antibody (ab290, Abcam) was crosslinked with magnetic Dynabeads Protein G (Invitrogen) at 0.5 μg/μl beads (50% slurry) with bis (sulfosuccinimidyl) suberate following the manufacturer’s instructions. The supernatant was then incubated with 25 μl of antibody-beads (50% slurry) at 4°C for 2 h. After immunoprecipitation, beads were washed with wash buffer (50 mM Tris-HCl, 150 mM NaCl, 0.1% Triton X-100), and two more times with wash buffer excluding the detergent (50 mM Tris-HCl, 150 mM NaCl).

### Protein digestion and nano LC-MS/MS data acquisition

The immunoprecipitated beads were suspended in 20 μl of digestion buffer (8 M urea, 250 mM Tris-HCl [pH 8.5]), reduced with 25 mM tris(2-carboxyethyl)phosphine (TCEP) at 37°C for 15 min and alkylated using 25 mM iodoacetamide at 37°C for 30 min in the dark, both with shaking at 1,200 rpm following the standard protocol. Proteins on beads were directly digested with 0.1 μg Lys-C (FUJIFILM Wako, Japan) at 37°C for 3 h with shaking at 1,000 rpm. After dilution to a urea concentration of 2 M with 50 mM Tris-HCl (pH 8.5) followed by addition of 1 mM CaCl_2_, Lys-C digest was further digested with 0.1 μg trypsin (Promega) with shaking at 1,000 rpm at 37°C overnight. The digestion was stopped by adding 5 μl of 20% TFA, and then Lys-C/trypsin– double digested peptides were desalted using a GL-Tip SDB (GL Science) according to the manufacturer’s instructions. Nano-LC-MS/MS analysis was performed using a EASY-nLC 1200 LC system (ThermoFisher Scientific) connected to a Orbitrap Exploris 480 hybrid quadrupole-orbitrap mass spectrometer (Thermo Fisher Scientific). Desalted samples were dissolved in 200 μl of 2% acetonitrile (0.1% TFA) and 7.5-μl aliquots of the supernatant were used for LC-MS/MS analysis. Samples were loaded in direct injection mode and separated on a nano-HPLC capillary column (Aurora column [75 μm I.D. × 250 mm], IonOpticks) with a gradient of 3–32% acetonitrile (containing 0.1% formic acid) over 90 min at a flow rate of 300 nl/min. The Orbitrap Exploris 480 mass spectrometer was operated in data-dependent acquisition mode with dynamic exclusion enabled (20 s). MS/MS scans were performed by higher-energy collisional dissociation with the normalized collision energy set to 30. The MS/MS raw files were processed and analyzed with Proteome Discoverer 2.5 (Thermo Fisher Scientific) using the SEQUEST HT algorithm, searching against the TAIR10 *Arabidopsis* protein database. The volcano plots were generated based on the calculated protein abundance^32^. Any missing values were imputed using a random selection from the bottom 5% of all protein abundances.

### Yeast Two-Hybrid Assays

Yeast two-hybrid screening was performed using the Matchmaker Gold Yeast Two-Hybrid System (Takara). The cDNA fragments corresponding to CEPDL2 and GrxS5 were subcloned into the pGBKT7 vector, which was subsequently introduced into the Y2H Gold yeast strain. The cDNA fragments for TGA1 and TGA4 were subcloned into pGADT7 vector and transformed into the Y187 yeast strain. After the mating process between the Y2H Gold strain expressing CEPDL2 or GrxS5 and the Y187 strain expressing TGA1 or TGA4, resulting zygotes were selected by culturing on SD/−Leu/−Trp/−Ade/−His +X-α-Gal plates for 5 days at 28°C.

### Root imaging

Cell outlines were stained with 50 µg/ml propidium iodide for 2 min and observed under a confocal laser-scanning microscope (Olympus FV3000) with helium-neon laser excitation at 543 nm. GFP images were collected with argon laser excitation at 488 nm.

### ChIP assay

The roots of 16-day-old TGA1–GFP plants grown on 10 mM NO_3_^−^ medium (∼900 mg) were fixed in 30 mL of 1% formaldehyde under vacuum for 5 cycles of 3 min. Cross-linking was quenched in 2 M glycine under vacuum for 3 min. Quenched samples were washed twice with 30 ml of cold 1× PBS on ice, dried briefly on paper towels, frozen in liquid nitrogen and then stored at −80°C. Frozen samples were ground to a fine powder using a Multi-beads shocker and dissolved in 3.5 mL of nuclei extraction buffer (10 mM Tris-HCl [pH 8.0], 0.25 M sucrose, 10 mM MgCl_2_, 40 mM β-mercaptoethanol, and protease inhibitor cocktail). Samples were filtered through two layers of Miracloth (Calbiochem, CA, USA) and centrifuged for 5 min at 11,000 rpm at 4°C. The pellets were resuspended in 75 μL of nuclei lysis buffer (50 mM Tris-HCl [pH 8.0], 10 mM EDTA, 1% SDS). After incubation on ice for 30 min, samples were mixed with 625 μL of ChIP dilution buffer without Triton (16.7 mM Tris-HCl [pH 8.0], 167 mM NaCl, 1.2 mM EDTA, and 0.01% SDS). Chromatin samples were sonicated for 100 cycles of 15 s ON/45 s OFF using a Bioruptor UCD-250 (Cosmo Bio Co., Ltd.) to produce DNA fragments, followed by the addition of 200 μL of ChIP dilution buffer with Triton (16.7 mM Tris-HCl [pH 8.0], 167 mM NaCl, 1.2 mM EDTA, 0.01% SDS, and 1.1% Triton X-100). After twice centrifugations for 5 min at 15,000 rpm at 4°C, the supernatant was collected in a new tube. 900 μL of solubilized sample was incubated with 25 μl of Anti-GFP antibody or Rabbit IgG-Isotype Control crosslinked beads at 4 °C for 16 h while an 18 μL aliquot was used as the input control. Beads were washed twice with 1 mL of low salt wash buffer (20 mM Tris-HCl [pH 8.0], 150 mM NaCl, 2 mM EDTA, 0.1% SDS, and 1% Triton X-100), twice with 1 mL of high salt wash buffer (20 mM Tris-HCl [pH 8.0], 500 mM NaCl, 2 mM EDTA, 0.1% SDS, and 1% Triton X-100), twice with 1 mL of LiCl wash buffer (10 mM Tris-HCl [pH 8.0], 0.25 M LiCl, 1 mM EDTA, 1% sodium deoxycholate, and 1% Nonidet P-40), and twice with 1 mL of TE buffer (10 mM Tris-HCl [pH 8.0], 1 mM EDTA). After washing, beads were resuspended in 100 μL of elution buffer (50 mM Tris-HCl [pH 8.0], 10 mM EDTA, 1% SDS) and incubated at 65°C for 30 min. 82 μL of elution buffer was added to 18 μL of the input control sample. Both supernatant and input samples were mixed with 6 μL of 5 M NaCl and incubated at 65°C overnight to reverse crosslinks. ChIP samples were mixed with 550 μL of PB Buffer and purified using the QIAquick PCR Purification Kit (Qiagen) following the manufacturer’s instructions.

### Construction of Illumina sequencing libraries and sequencing of ChIP DNA

ChIP-seq libraries were constructed from 100 ng of DNA samples using the NEB Ultra II DNA Library Prep Kit for Illumina (New England BioLabs) according to the manufacturer’s instructions. The amount of cDNA was determined by the QuantiFluor dsDNA System (Promega, WI, USA). All ChIP-seq libraries were sequenced as 81-bp single-end reads using the Illumina system NextSeq550.

### Analysis of ChIP-Seq data

Reads were mapped to the TAIR10 *Arabidopsis* genome database using Bowtie2 with default parameters^33^. The Sequence Alignment/Map (SAM) file generated by Bowtie2 was converted to a Binary Alignment/Map (BAM) format file by SAMtools^34^. To visualize the mapped reads, the Tiled Data File (TDF) file was generated from the BAM file using the igvtools package in the Integrative Genome Browser (IGV)^35^. ChIP-seq peaks were called by comparing the IP (Anti-GFP) with the Input (Rabbit IgG-Isotype Control) using Model-based Analysis of ChIP-Seq (MACS2) with the “-q 0.01” option (q value < 0.01)^36^. The peaks were annotated with the nearest gene using the Bioconductor and ChIPpeakAnno package in R program^37, 38^. Gene Ontology (GO) analysis of the set of 1,105 genes was performed by PANTHER^39^ in The *Arabidopsis* Information Resource (TAIR) website (https://www.arabidopsis.org/tools/go_term_enrichment.jsp). Sequences of the peaks were extracted from the *Arabidopsis thaliana* genome as a FASTA file by Bedtools^40^. To identify the candidates of NPR1 binding motifs, the FASTA files were subjected to MEME (Multiple EM for Motif Elicitation)-ChIP with the default parameter (–meme-minw 6–meme-maxw 10)^41^ and generated a density plot of the distribution of the motifs.

### ChIP-qPCR analysis

The roots of 16-day-old *TGA1–GFP*/Nössen and *TGA1–GFP*/*cepd1,2 cepdl2* plants grown on 10 mM NO_3_^−^ medium (∼900 mg) were used for ChIP-qPCR. Chromatin samples were sonicated for 30 cycles of 15 s ON/45 s OFF using a Bioruptor UCD-250 to obtain longer DNA fragments than ChIP-seq analysis. Real-time RT-qPCR was carried out on a StepOne System (Applied Biosystems) using the THUNDERBIRD Next SYBR qPCR Mix (Toyobo), 10-fold diluted purified DNAs, and gene-specific primers.

### Real-time RT-qPCR

Total RNA was obtained from roots or shoots using an RNeasy kit (Qiagen). First-strand cDNA was synthesized from 0.5 μg of total RNA using the Superscript IV VILO Master Mix (Thermo Fisher Scientific) according to the manufacturer’s protocol. Primers and Taqman probes were designed using Probe Finder software in the Universal Probe Library (UPL) Assay Design Center (Roche). All PCR reactions were performed using a StepOne System (Applied Biosystems). Constitutively expressed EF-1α was used as a reference gene for normalization of quantitative RT-PCR data.

### Promoter GUS analysis

For β-glucuronidase (GUS) reporter-aided analysis of *GrxS1-S8* promoter activity, we amplified the upstream 2.0-kb promoter regions of the genes using genomic PCR and cloned them by translational fusion in frame with the β-glucuronidase (GUS) coding sequence in the binary vector pBI101. GUS activity was visualized using a standard protocol with X-Gluc as the substrate. For leaf sectioning, leaves were fixed in FAA solution (3.7% formaldehyde, 5% acetic acid and 50% ethanol), dehydrated through a graded ethanol series and embedded in Technovit 7100 resin (Heraeus Kulzer, Germany) following the manufacturer’s protocol. Sections were cut at 5 μm thickness using a rotary microtome (Leica RM2235), counter-stained with 0.05% Nile red, mounted with Entellan (Merck) and observed under a standard light microscope (Olympus BX60).

### Nitrate content

The concentration of NO_3_^−^ ions in tissues was determined using an ion chromatography system (Dionex Aquion, Thermo Fisher Scientific). Shoots of 14-day-old plants were powdered in liquid nitrogen and mixed with 1 ml of water to extract nitrate. After centrifugation, the crude tissue extract was diluted 10-fold with water, and 25-μl aliquots were analyzed on a Dionex IonPac AS22 column (4 mm i.d. × 250 mm) over 15 min. The mobile phase eluent consisted of 4.5 mM Na_2_CO_3_ and 1.4 mM NaHCO_3_, employed at a flow rate of 1.2 ml/min at 30°C, and separation was monitored using a conductivity detector equipped with a Dionex AERS 500 suppressor unit.

### Root ^15^N influx

Vertically grown 14-day-old plants were sequentially transferred to 0.1 mM CaSO_4_ for 1 min and then to modified Murashige-Skoog medium containing 0.2 mM or 10 mM ^15^NO_3_^−^ as the N source for 10 min. At the end of ^15^N labeling, the roots were washed for 1 min in 0.1 mM CaSO_4_ and separated from the shoots. The roots were lyophilized *in vacuo* and analyzed for total N and ^15^N content using elemental analysis–isotope ratio mass spectrometry (Flash EA1112-DELTA V PLUS ConFlo III system, Thermo Fisher Scientific).

### RNA-seq analysis

Total RNA was extracted from roots of a 14-day-old Col and *GFP-GrxS5* plants grown on 10 mM NO_3_^-^ using RNeasy Mini Kit (QIAGEN). We used 1 μg total RNA for mRNA purification using the NEBNext Oligo d (T) 25 magnetic isolation module (New England Biolabs), followed by first strand cDNA synthesis using NEBNext Ultra II RNA Library Prep Kit for Illumina according to the manufacturer’s protocols. Samples were then ligated with NEBNext multiplex oligo adaptor kits for barcoding. The amount of cDNA was determined on an Agilent 4150 TapeStation System. The resulting cDNA libraries were sequenced on an Illumina NextSeq 550 with single-end 81-bp sequencing, respectively. The reads were mapped to the Arabidopsis TAIR10 reference genome using BaseSpace software on the web (Illumina, https://basespace.illumina.com). Pairwise comparisons between samples were performed with the EdgeR package on the web (Degust, https://degust.erc.monash.edu)^42^. Genes with a *q* value < 0.05 and absolute fold change < 0.7 were defined as differentially expressed genes. Data shown are the mean of at least three biologically independent RNA-seq data.

### Hydroponic culture system

In a hydroponic system, we used modified Murashige-Skoog liquid medium without sucrose to prevent bacterial growth. Thirteen-day-old *Arabidopsis* seedlings were transplanted into a hydroponic system. Individual roots were carefully guided through pre-drilled apertures (5 mm I.D.) in a Styrofoam plate, and subsequently the plate along with the plants was positioned on a 6-well culture plate filled with 1 mM NO_3_^−^ liquid medium. The plant roots were thoroughly immersed in the medium and subjected to a pre-incubation for 24 h. WT and *tga1,4* plants were cultured under continuous nitrogen-sufficient (1 mM NO_3_^−^), continuous low nitrogen (0.1 mM NO_3_^−^) or fluctuating nitrogen condition where 18 h of nitrogen-deficient (0 mM NO_3_^−^) and 6 h of nitrogen-sufficient (1 mM NO_3_^−^) periods were alternated for 7 d.

### Stable isotope metabolic labeling with ^14^N or ^15^N

For quantitative proteomic analyses, WT (Nössen) and *tga1,4* (Nos) seedlings were reciprocally labeled with light (normal) ^14^N or heavy ^15^N via metabolic incorporation. We defined ^14^N-labeled WT/^15^N-labeled *tga1,4* pairs as forward labeling and ^15^N WT/^14^N *tga1,4* pairs as reciprocal labeling. Surface-sterilized WT and *tga1,4* seeds were sown on either ^14^N- or ^15^N-containing B5 medium (1% sucrose) solidified using 1.5% agar in 13 × 10 cm plastic plates (18 seeds × 3 row/plate) and then grown vertically at 22°C for 7 d with continuous light. In ^15^N B5 medium, K^14^NO_3_ and (^14^NH_4_)_2_SO_4_ were substituted with heavy nitrogen salts, K^15^NO_3_ and (^15^NH_4_)_2_SO_4_, respectively. After 7 d, the seedlings were transferred to 3 mM NO_3_^−^ medium (with either K^14^NO_3_ or K^15^NO_3_) solidified using 1.5% purified agar in 13 × 10 cm plastic plates (12 seedlings/plate) and further grown vertically at 22°C for 7 d with continuous light. And then, these plants were transplanted into a hydroponic system where 18 h of N-deficient (0 mM NO_3_^−^) and 6 h of N-sufficient (1 mM NO_3_^−^) periods were alternated for 2 d. For forward labeled samples, ^14^N-labeled WT roots were combined at a 1:1 fresh weight ratio with ^15^N-labeled *tga1,4* roots. Similarly, ^15^N-labeled WT roots were combined at a 1:1 ratio with ^14^N-labeled *tga1,4* roots for reciprocal labeling. The combined root tissues (0.5-1.0 g) were frozen in liquid nitrogen and ground to a fine powder using a Multi-beads shocker. Microsomal membranes were extracted by mixing 500 mg of ground tissue with 2.5 ml of extraction buffer (25 mM Tris-HCl [pH 7.0], 10 mM MgCl_2_, 2 mM DTT, 2 µM leupeptin, 2 mM PMSF, 250 mM sucrose, and 1× PhosSTOP phosphatase inhibitors [Roche]). The homogenate was centrifuged for 15 min at 10,000 × *g* at 4°C, and then the supernatant was further centrifuged at 100,000 g for 30 min at 4 °C to give a pellet of microsomal membranes. The pellets were solubilized in digestion buffer (8 M urea, 250 mM Tris-HCl [pH8.5]) at a final protein concentration of 2.0 mg/ml. The protein concentration was determined using a Bradford protein assay kit. Protein digestion and nano LC-MS/MS data acquisition method are described above.

## Data Availability Statement

The RNA-seq data have been deposited in the NCBI Gene Expression Omnibus (GEO) under accession number GSE240738. The raw mass spectrometry data have been deposited in the ProteomeXchange Consortium via PRIDE partner repository under accession number PXD044498. The source data underlying Figs. 1–5 and Supplementary Figs. 1–3 are provided as a Source Data file. The *Arabidopsis* lines generated in this study are available from the corresponding author on reasonable request.

## Acknowledgments

This research was supported by a Grant-in-Aid for Scientific Research (S) (No. 23H05477 to Y.M.), Grant-in-Aid for Transformative Research Areas (A) (No. 20H05907 to Y.M.) and a Grant-in-Aid for JSPS Fellows (No. 22KJ1616 to Y.O.) from the Japan Society for the Promotion of Science.

## Author Contributions

Y.M. conceived this project and Y.M. R.K. Y.O. and M.O.-O. designed the experiments. All authors performed the experiments and interpreted the results. Y.M. and Y.O. wrote the manuscript.

---

**Supplementary Fig. 1.**
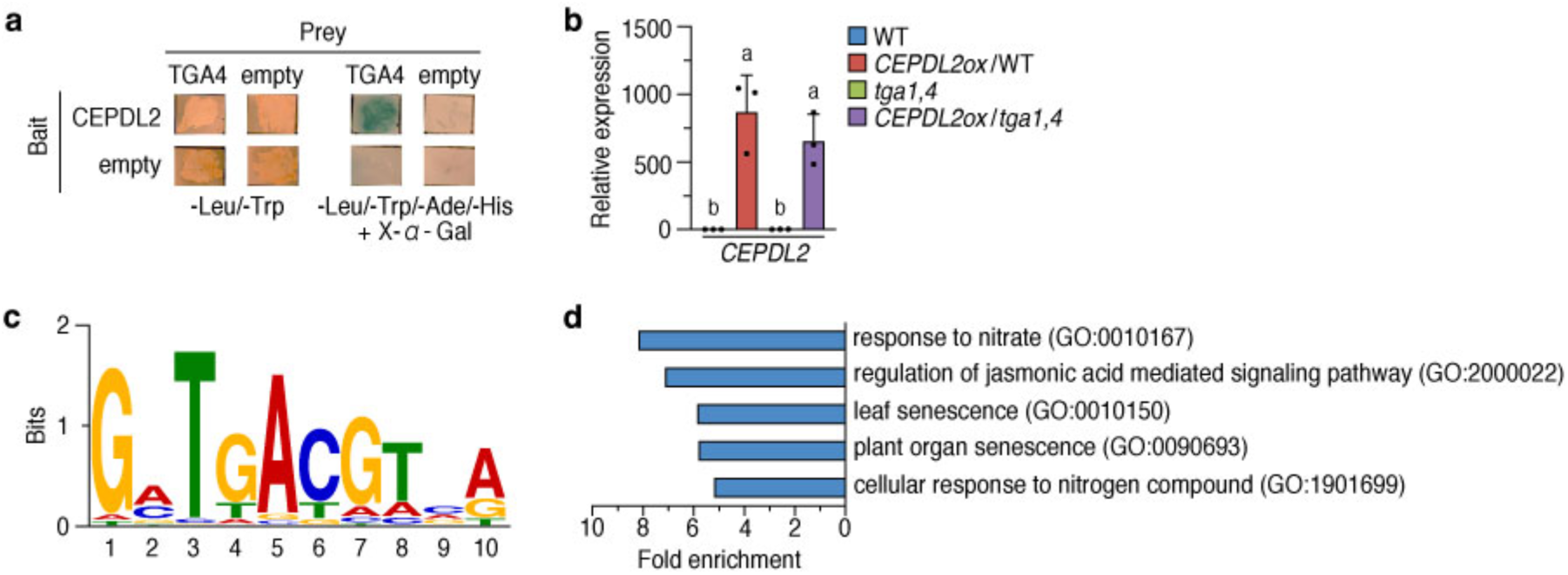
*TGA1* and *TGA4* acts downstream of *CEPDL2*. (**a**) Yeast two-hybrid assays show the interaction between CEPDL2 and TGA4. (**b**) Overexpression levels of *CEPDL2* in each transgenic line (mean ± SD, *P < 0.05, *n* = 3). (**c**) Binding motifs for TGA1 enriched in the ChIP-Seq reads. (**d**) Bar plot showing the top 5 enriched GO terms.

**Supplementary Fig. 2.**
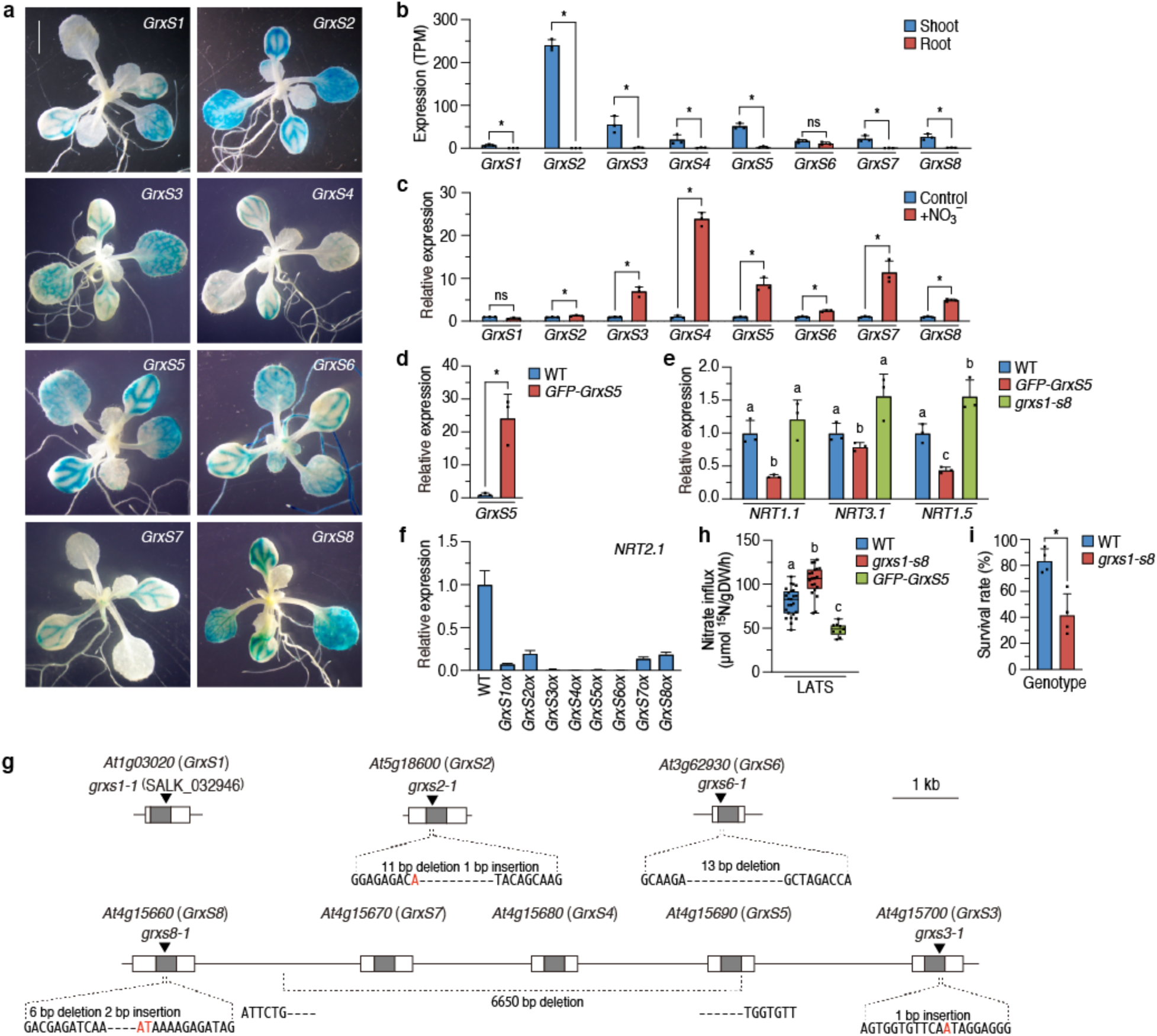
GrxS1-S8 family polypeptides repress nitrate uptake genes. (**a**) Histochemical staining for GUS activity in 10-day-old transgenic plants expressing the *GUS* gene under control of each *GrxS* promoter. Scale bar = 5 mm. (**b**) Expression levels of *GrxS1* to *GrxS8* genes in shoots and roots of WT plants (mean ± SD, *P < 0.05, *n* = 3). (**c**) qRT-PCR analysis of *GrxS1* to *GrxS8* transcripts in the detached leaves of 14-day-old WT and plants after 10 mM nitrate treatment for 3 h (mean ± SD, *P < 0.05, *n* = 3). (**d**) Expression level of *GrxS5* in the transgenic plants expressing *GFP-GrxS5* (mean ± SD, *P < 0.05, *n* = 3). (**e**) Expression levels of nitrate uptake genes in WT, *GFP-GrxS5* and *grxs1-s8* plants grown on 10 mM NO_3_^−^ medium (mean ± SD, P < 0.05, *n* = 3). (**f**) Expression level of *NRT2.1* in the transgenic plants overexpressing *GrxS1* to *GrxS8* (mean ± SD). (**g**) Schematic representation of the deletion sites in octuple mutant *grxs1-s8*. Nucleotide insertions are shown in red. (**h**) Activity of low-affinity nitrate uptake system (LATS) of 14-day-old WT, *grxs1-s8* and *GFP-GrxS5* plants grown on 10 mM NO_3_^−^ medium (mean ± SD, P < 0.05, *n* = 9-25). (**i**) Survival rate of WT and *grxs1-s8* octuple mutant under high nitrate (120 mM) treatment (mean ± SD, *P < 0.05, *n* = 4).

**Supplementary Fig. 3.**
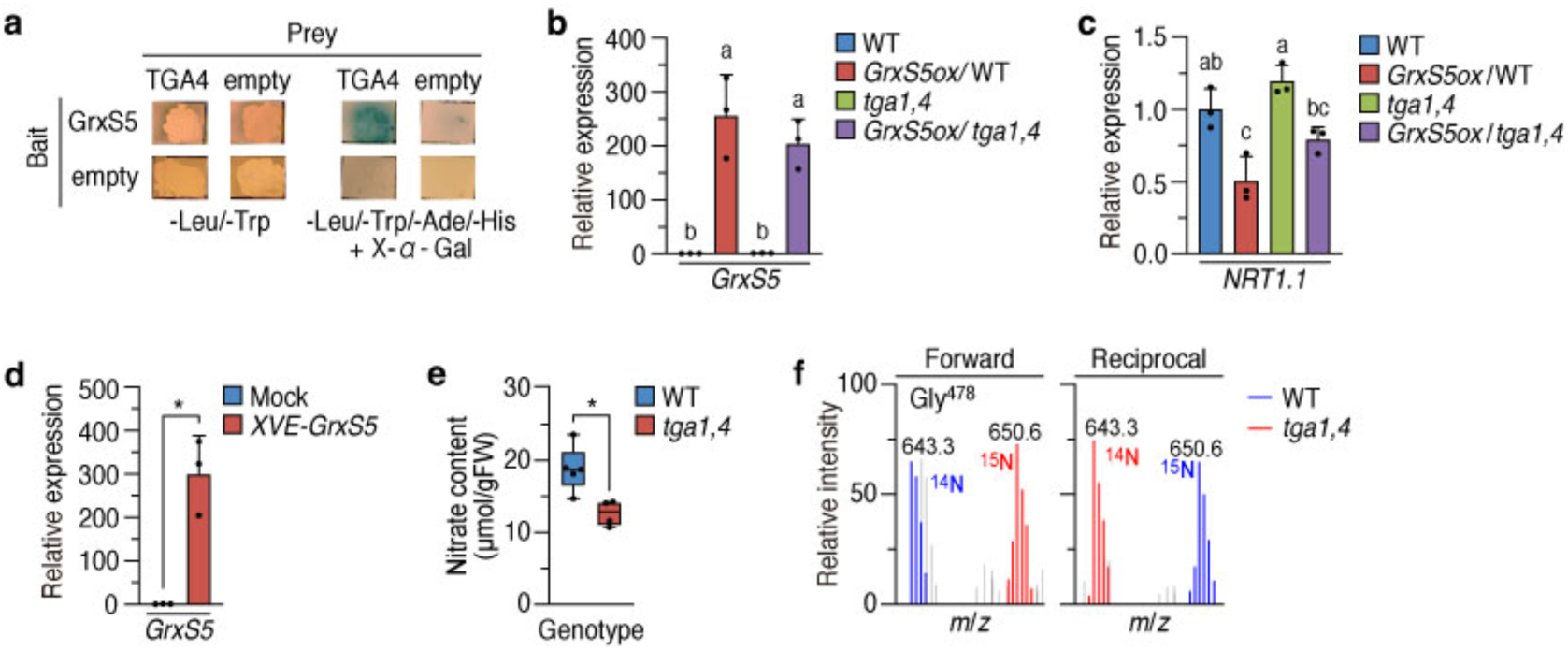
CEPDL2 and GrxS5 competitively target TGA1/4. (**a**) Yeast two-hybrid assays show the interaction between GrxS5 and TGA4. (**b**) Overexpression levels of *GrxS5* in each transgenic line. (**c**) Expression level of *NRT1.1* in WT, *GrxS5ox*/WT, *tga1,4* and *GrxS5ox*/*tga1,4* plants grown on 10 mM NO_3_^−^ medium (mean ± SD, P < 0.05, *n* = 3). (**d**) Overexpression levels of *GrxS5* in XVE-GrxS5 plants (mean ± SD, *P < 0.05, *n* = 3). (**e**) Nitrate content in shoots of WT and *tga1,4* plants grown for 72 h under fluctuating nitrogen condition where 18 h of nitrogen-deficient (0 mM NO_3_^−^) and 6 h of nitrogen-sufficient (1 mM NO_3_^−^) periods were alternated. (**f**) Mass spectra of the NRT2.1 Gly^478^ peptide (position 478-493 in NRT2.1 protein) derived from reciprocally ^14^N- and ^15^N-labeled plants, showing comparable NRT2.1 protein abundance in WT and *tga1,4* plants.

